# Kv1.1 subunits localize to cardiorespiratory brain networks in mice where their absence induces astrogliosis and microgliosis

**DOI:** 10.1101/2021.02.01.429209

**Authors:** Hemangini A Dhaibar, Kathryn A Hamilton, Edward Glasscock

**Affiliations:** Department of Cellular Biology & Anatomy, Louisiana State University Health Sciences Center, Shreveport, LA USA; Department of Biological Sciences, Southern Methodist University, Dallas, TX USA

**Author notes:** **Corresponding author:** Edward Glasscock, Department of Biological Sciences, Southern Methodist University, Dallas, TX.

## Abstract

Cardiorespiratory collapse following a seizure is a suspected cause of sudden unexpected death in epilepsy (SUDEP), the leading cause of epilepsy-related mortality. In the commonly used *Kcna1* gene knockout (*Kcna1^−/−^*) mouse model of SUDEP, cardiorespiratory profiling reveals an array of aberrant breathing patterns that could contribute to risk of seizure-related mortality. However, the brain structures mediating these respiratory abnormalities remain unknown. We hypothesize that Kv1.1 deficiency in respiratory control centers of the brain contribute to respiratory dysfunction in *Kcna1^−/−^* mice leading to increased SUDEP risk. Thus, in this study, we first used immunohistochemistry to map expression of Kv1.1 protein in cardiorespiratory brain regions of wild-type *Kcna1^+/+^* (WT) mice. Next, GFAP and Iba1 immunostaining was used to test for the presence of astrogliosis and microgliosis, respectively, in the cardiorespiratory centers of *Kcna1^−/−^* mice, which could be indicative of seizure-related brain injury that could impair breathing. In WT type mice, we detected Kv1.1 protein in all cardiorespiratory centers examined, including the basolateral amygdala, dorsal respiratory group, dorsal motor nucleus of vagus, nucleus ambiguus, ventral respiratory column, and pontine respiratory group, as well as chemosensory centers including the retrotrapezoid and median raphae nuclei. Extensive gliosis was observed in the same areas in *Kcna1^−/−^* mice suggesting that seizure-associated brain injury could contribute to respiratory abnormalities.

## 1. INTRODUCTION

Cardiorespiratory dysfunction is suspected as a primary risk factor for sudden unexpected death in epilepsy (SUDEP), the leading cause of epilepsy-related mortality, which is generally defined as the sudden death of someone with epilepsy for unknown reasons even following autopsy (Kennedy and Seyal 2015). Clinical studies of mortality in epilepsy monitoring units suggest that patients who succumb to SUDEP show a lethal cascade of seizure followed by terminal apnea, which appears to induce cardiac arrest (Ryvlin *et al*. 2013). Several mouse models also exhibit seizure-related respiratory arrest associated with increased risk of death, including the *Kcna1* gene knockout (*Kcna1^−/−^*) mouse which is commonly used to study pathomechanisms underlying SUDEP (Feng and Faingold 2017; Kim *et al*. 2018; Kruse *et al*. 2019; Glasscock *et al*. 2010; Dhaibar *et al*. 2019). *Kcna1^−/−^* mice lack Kv1.1 voltage-gated a-subunits, which control action potential firing properties in brain and heart, leading to spontaneous seizures that culminate in sudden premature death with onset at about two weeks old (Smart *et al*. 1998; Glasscock 2019). Cardiorespiratory profiling reveals primary breathing dysfunction during seizures in *Kcna1^−/−^* mice, including a high incidence of hyperventilation, tachypnea, and ataxic breathing, as well as occasional hypopnea, bradypnea, and apnea (Dhaibar *et al*. 2019). In addition, *Kcna1^−/−^* mice also exhibit basal breathing irregularities, including mild tachypnea, an absence of post-sigh apneas, a 3-fold increase in respiratory variability, and frequent hypoxia, which suggest Kv1.1 subunits play a fundamental role in regulating normal respiratory physiology (Dhaibar *et al*. 2019; Simeone *et al*. 2018).

The pathways and mechanisms contributing to breathing dysfunction in *Kcna1^−/−^* mice have not been identified in detail. Neuron–specific ablation of Kv1.1 in the central nervous system recapitulates most of the respiratory phenotypes seen in *Kcna1^−/−^* mice demonstrating a neural basis for the breathing abnormalities (Trosclair et al. 2020). In addition, Kv1.1 transcripts and protein are not detectable in the lungs of wild-type (WT) mice (Simeone *et al*. 2018). Kv1.1 exhibits widespread and primarily axonal expression in forebrain and cerebellar regions, but its localization in brainstem nuclei that control respiration has not been well characterized (Wang *et al*. 1994). In addition to the absence of Kv1.1 subunits in neurons that normally express them, Kv1.1 deficiency could also indirectly impair neural function by causing seizures which induce brain damage. One of the most common brain pathologies associated with epilepsy is reactive gliosis, a process whereby astrocytes and microglia become activated in response to injury and undergo morphological changes and increased proliferation that can promote neuronal hyperexcitability (Patel *et al*. 2019; Devinsky *et al*. 2013). In *Kcna1^−/−^* mice, seizures induce significant gliosis in the hippocampal formation and the hypothalamus, but other areas have not been investigated in detail (Wenzel *et al*. 2007; Roundtree *et al*. 2016).

Therefore, the goal of this work was to investigate the localization of Kv1.1 subunits in cardiorespiratory brain regions of WT mice and to determine whether the absence of Kv1.1 in *Kcna1^−/−^* mice leads to seizure–associated pathology in these neuronal populations that could potentially impair function. First, immunohistochemistry was used to map Kv1.1 protein localization in WT mice in the major brainstem nuclei that are responsible for the control of respiration and cardiac function. To identify seizure-related damage in cardiorespiratory centers, astrogliosis and microgliosis were then assessed in *Kcna1^−/−^* brains by imaging immunoreactivity for glial fibrillary acidic (GFAP) protein and for ionized calcium binding adaptor molecule 1 (Iba1) protein, respectively. Our findings reveal prominent Kv1.1 immunoreactivity in respiratory brainstem regions where the channels could influence breathing intrinsically, as well as extensive gliosis in respiratory nuclei in *Kcna1^−/−^* mice suggesting the potential for impaired breathing due to seizure-induced injury.

## 2. MATERIALS AND METHODS

### 2.1 Animals and genotyping

*Kcna1^−/−^* mice (Tac:N:NIHS–BC genetic background) carry null knockout (KO) alleles of the *Kcna1* gene generated by targeted deletion of the entire open reading frame on chromosome 6 (Smart *et al*. 1998). Mice were housed at 22°C, fed ad libitum, and submitted to a 12-h light/dark cycle. For genotyping, genomic DNA was isolated by enzymatic digestion of tail clips using Direct-PCR Lysis Reagent (Viagen Biotech, Los Angeles, CA, USA). Genotypes were determined by performing PCR amplification of genomic DNA using allele-specific primers. The following primer sequences were used to yield genotype–specific amplicons of ~337 bp for the wild-type (WT) allele and ~475 bp for the KO allele: a WT-specific primer (5’-GCCTCTGACAGTGACCTCAGC-3’), a KO-specific primer (5’-CCTTCTATCGCCTTCTTGACG-3’), and a common primer (5’-GCTTCAGGTTCGCCACTCCCC-3’). All experiments were performed in accordance with National Institutes of Health (NIH) guidelines with approval from the Institutional Animal Care and Use Committee of the Louisiana State University Health Sciences Center-Shreveport.

### 2.2 Immunohistochemistry

Age-and sex-matched 6-month old WT (*Kcna1^+/+^*; n=5: 3 males, 2 females) and KO mice (*Kcna1^−/−^*; n=4: 2 males, 2 females) were used for immunohistochemical analysis. Older mice were selected for study to examine the long-term effects of chronic seizures, which begin at about 2 weeks old, thereby increasing the probability of accumulated brain pathology. Mice were transcardially perfused, first with phosphate buffered saline (PBS), then with 4% paraformaldehyde in PBS and lastly with 4% sucrose in PBS. After perfusion, the brains were removed and cryoprotected at 4°C for 3 days in a series of ascending sucrose solutions (10%, 20%, and 30% sucrose in PBS for one day each). Brains were then embedded in optimum cutting temperature medium (OCT) and frozen by placing at −80°C for 1 h. Brains were kept at −20°C for ~2 h immediately before coronal sectioning (10 μm) using a cryostat maintained at −20°C. Nuclear and antibody staining of serial sections was used to identify the following anatomical regions that fall within the Bregma coordinates as shown in the mouse brain atlas (Franklin and Paxinos, 2013): basolateral amygdala (Bregma −0.80 to −0.90 mm); hippocampus (Bregma −1.50 to −1.70 mm); parabrachial nucleus (PBP; Bregma −3.30 to −3.90 mm); retrotrapezoid nucleus (RTN), median raphae nucleus (MnR) and Kölliker-Fuse nucleus (KF; Bregma −4.80 to −5.15 mm); cerebellum (Bregma −5.40 to −8.00 mm); nucleus solitary tract (NTS; Bregma −6.50 to −7.00 mm); Bötzinger complex (Bregma −6.60 to −6.75 mm); nucleus ambiguus (NA; Bregma −6.60 to −7.30 mm); preBötzinger complex (Bregma Bregma −6.83 to −7.10 mm); rostral ventral respiratory group (RVRG; Bregma −7.20 to −7.30 mm); and dorsal motor nucleus of vagus (10N) and hypoglossal nucleus (12N; Bregma −7.20 to −7.70 mm). After cutting, sections were blocked and permeabilized for 1 h in antibody vehicle (10% BSA, 0.3% Triton X-100 in PBS), followed by incubation in primary antibodies for 15-20 h at room temperature (~22 °C). Subsequently, sections were washed 3 times in vehicle and incubated for 1 h in secondary antibodies. Sections were then washed with vehicle once and PBS 3 times prior to staining with NucBlue Fixed Cell ReadyProbes reagent (Invitrogen; Life Technologies Co., Eugene, OR). Following a final wash in PBS, sections were slide-mounted using ProLong Glass Antifade Mountant without DAPI (Invitrogen; Life Technologies Co., Eugene, OR). Negative control experiments were performed by incubating sections with secondary antibodies in vehicle. Images were captured using an Olympus IX71 epifluorescence microscope with iVision as the acquisition software. Images were always acquired using the same optimized settings for each fluorophore. After acquisition, images were colorized using Fiji software, an image-processing package of ImageJ software (National Institutes of Health; Bethesda, MD, USA). Brightness and contrast were minimally adjusted to improve visualization of acquired images, but these settings were applied equally for each fluorophore for all images. Merged images were not further adjusted. Immunoreactivity was graded according to the approximate density of labeling and then assigned an incremental score of – (absent), + (low), ++ (moderate), or +++ (high).

### 2.3 Antibodies

The following primary antibodies and dilutions were used: mouse monoclonal anti-Kv1.1, 1:1000 (clone K20/78, Antibodies Incorporated, Davis, CA); rabbit polyclonal anti-neurokinin receptor 1 (NK1R), 1:5000 (S8305, Sigma-Aldrich, St. Louis, MO); rabbit polyclonal anti-Forkhead box protein P2 (FOXP2), 1:8000 (ab16046, Abcam, Cambridge, United Kingdom); mouse monoclonal anti-glial fibrillary acidic protein (GFAP) supernatant, 1:20 (clone N206A/8, Antibodies Incorporated, Davis, CA); rabbit polyclonal anti-ionized calcium binding adaptor molecule 1 (Iba1), 1:1000 (019–19741, Wako Chemicals, Richmond, VA). Secondary antibodies included Alexa Fluor 488 goat anti-mouse IgG1, 1:1000 (Invitrogen; Carlsbad, CA) and Alexa Fluor 594 goat anti-rabbit IgG, 1:1000 (Invitrogen; Carlsbad, CA).

## 3. RESULTS

### 3.1 Distribution of Kv1.1 immunoreactivity in wild-type (WT) mouse brain

Limbic system structures including the amygdala and hippocampus contain neurons that modulate respiration (Homma and Masaoka 2008; Dlouhy *et al*. 2015; Lhatoo *et al*. 2015; Nobis *et al*. 2019). Kv1.1 protein expression patterns have already been described in detail in hippocampus but not in the basolateral amygdala, which is known to be strongly activated by seizures in KO mice (Wang *et al*. 1994; Prüss *et al*. 2010; Gautier and Glasscock 2015). Moderate Kv1.1 immunoreactivity was detected in neuropil and some cell somata in basolateral amygdala (**Figure 1**). The density of Kv1.1 immunoreactivity in the amygdala and other brain regions examined is summarized in **Table 1**. Importantly, KO brains exhibited no significant Kv1.1 immunoreactivity in any brain regions demonstrating the specificity of the Kv1.1 antibody staining.

**Figure 1.**
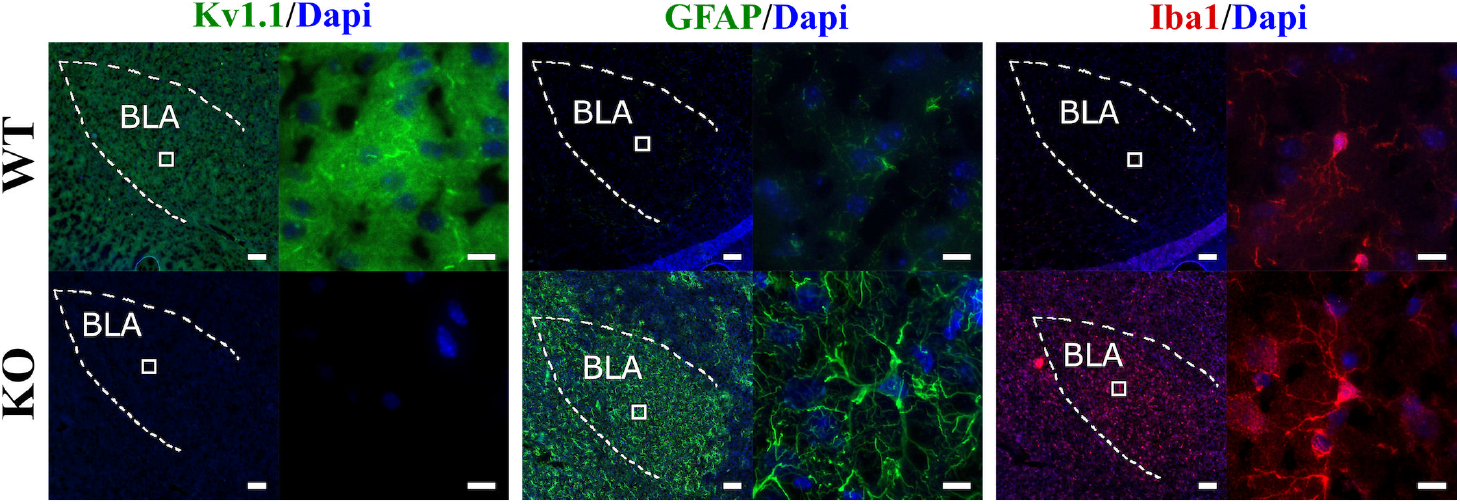
Kv1.1, GFAP, and Iba1 immunoreactivity in basolateral amygdala (BLA) of wild–type (WT) and *Kcna1^−/−^* (KO) mice. In each figure, the left panel shows a lower magnification image (scale bar = 100 μm) and the right panel shows a higher magnification image (scale bar = 10 μm) of the boxed region. The dotted lines represent the boundary of the brain region. DAPI staining was used to label the nuclei of cells.

**Table 1.**
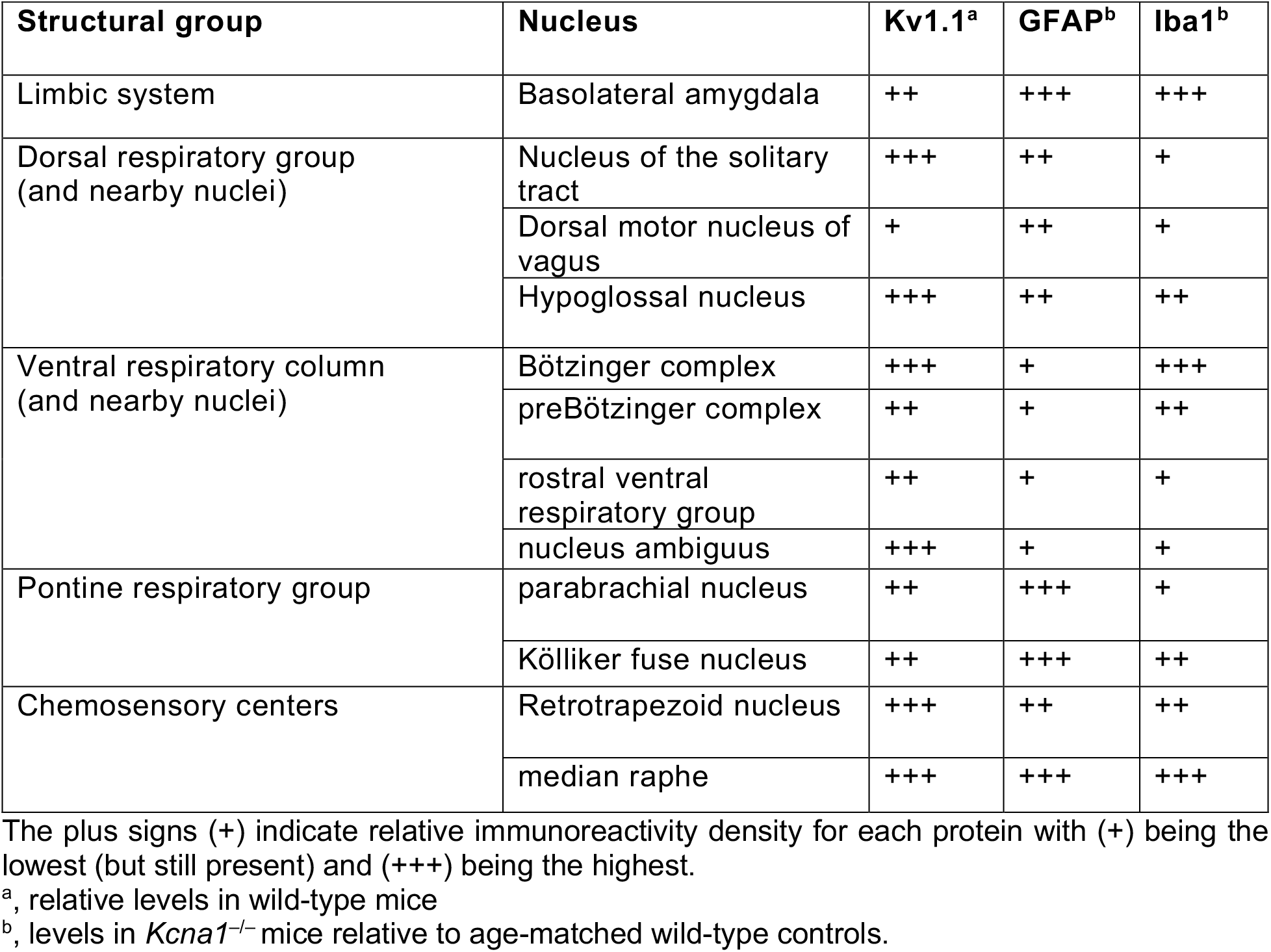
Summary of Kv1.1, GFAP, and Iba1 immunoreactivity density in mouse respiratory brain regions.

| Structural group | Nucleus | Kv1.1<sup>a</sup> | GFAP<sup>b</sup> | Iba1<sup>b</sup> |
| --- | --- | --- | --- | --- |
| Limbic system | Basolateral amygdala | ++ | +++ | +++ |
| Dorsal respiratory group<br>(and nearby nuclei) | Nucleus of the solitary tract | +++ | ++ | + |
|  | Dorsal motor nucleus of vagus | + | ++ | + |
|  | Hypoglossal nucleus | +++ | ++ | ++ |
| Ventral respiratory column<br>(and nearby nuclei) | Bötzinger complex | +++ | + | +++ |
|  | preBötzinger complex | ++ | + | ++ |
|  | rostral ventral respiratory group | ++ | + | + |
|  | nucleus ambiguus | +++ | + | + |
| Pontine respiratory group | parabrachial nucleus | ++ | +++ | + |
|  | Kölliker fuse nucleus | ++ | +++ | ++ |
| Chemosensory centers | Retrotrapezoid nucleus | +++ | ++ | ++ |
|  | median raphe | +++ | +++ | +++ |

The plus signs (+) indicate relative immunoreactivity density for each protein with (+) being the lowest (but still present) and (+++) being the highest.
<sup>a</sup>, relative levels in wild-type mice
<sup>b</sup>, levels in *Kcna1*<sup>-/-</sup> mice relative to age-matched wild-type controls.

The nucleus of the solitary tract (NTS) contains clusters of respiratory-modulated inspiratory neurons that act to coordinate respiratory reflex responses to afferent cardiorespiratory inputs from peripheral chemoreceptors, baroreceptors, and pulmonary stretch receptors (Zoccal *et al*. 2014). In NTS, neuropil and paired punctae consistent with juxtaparanodeal immunoreactivity of myelinated axons was observed, as previously observed for the vagus nerve (Glasscock *et al*. 2012; Glasscock *et al*. 2010; Wang *et al*. 1994) (**Figure 2a**). The localization of the axons is similar to the path of axons from the baroreceptive region of the NTS (Li *et al*. 2016). Thus, NTS, which is known for an important role in baroreception and respiration, exhibits significant Kv1.1 staining.

**Figure 2.**
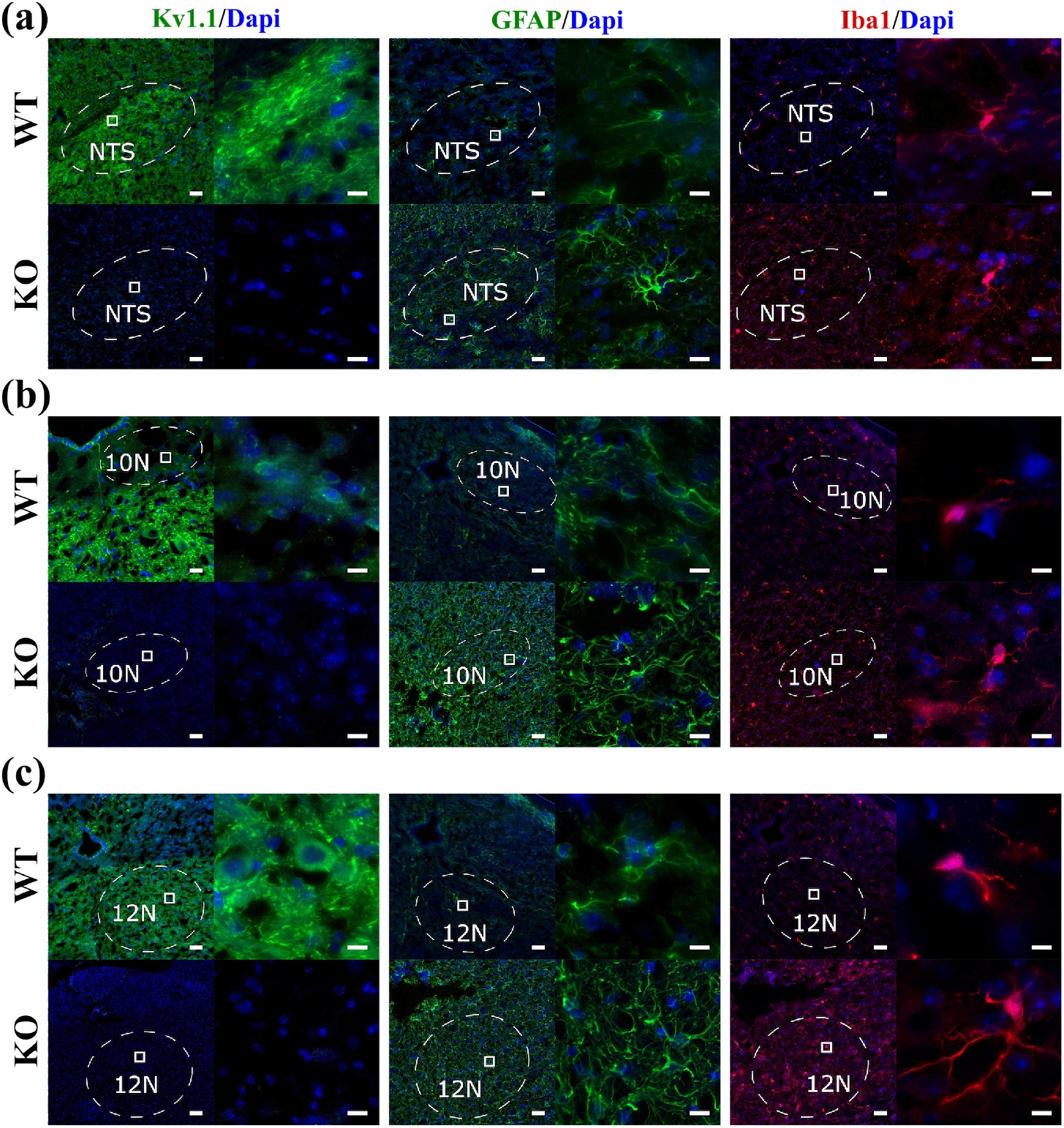
Kv1.1, GFAP, and Iba1 immunoreactivity in the dorsal respiratory group and nearby cardiorespitory nuclei in wild–type (WT) and *Kcna1^−/−^* (KO) mice. The structures shown include the (a) nucleus of the solitary tract (NTS), (b) dorsal motor nucleus of vagus (10N), and (c) hypoglossal nucleus (12N). For each condition, the left panel shows a lower magnification image (scale bar = 100 μm) and the right panel shows a higher magnification image (scale bar = 10 μm) corresponding to the boxed region in the left panel. The dotted lines represent the boundary of the respective brain region being examined. DAPI staining was used to label the nuclei of cells.

The dorsal motor nucleus of vagus (10N) is located along the floor of the fourth ventricle in the medulla and it is the nucleus for the vagus nerve, which provides parasympathetic efferents to the heart and other organs (Thompson *et al*. 2019). No Kv1.1 immunoreactivity was observed in 10N at lower magnification, but at higher magnification, low density Kv1.1 staining could be seen in some cell bodies (**Figure 2b**).

The hypoglossal nucleus (12N) is a cranial nerve motor nucleus which contains neurons that exhibit pre-inspiratory and inspiratory discharges that stabilize the upper airway prior to the commencement of airflow to the lungs (St.-John *et al*. 2004). Reduction in eupneic ventilatory drive is observed when these discharges are uncoupled (St.-John *et al*. 2004). Kv1.1 was highly expressed in the somata of these neurons and in associated fiber tracts (**Figure 2c**).

The ventral respiratory column (VRC) includes the Bötzinger nucleus, preBötzinger complex, and the rostral ventral respiratory group (RVRG) (Smith *et al*. 2009). The neurons of the VRC are involved in respiratory rhythm generation and their activity modulates the amplitude of respiratory motor output to spinal respiratory nerves (Alheid and McCrimmon 2008). The neighboring nucleus ambiguus (NA) houses preganglionic parasympathetic vagal neurons that innervate the heart and muscles of the soft palate, larynx, pharynx, adjusting ventilatory drive to respiratory muscle activity (Wijdicks 2017; Thompson *et al*. 2019). To aid in identification of the VRC and NA, Neurokinin–1 (NK1R) protein immunoreactivity was used as a marker since it specifically labels neurons of the Bötzinger nucleus, pre-Bötzinger nucleus, RVRG, and NA (Forsberg *et al*. 2016; Wang *et al*. 2002). Moderate to high density somatic Kv1.1 staining was observed in cells of all VRC nuclei examined (**Figure 3a-c**). Kv1.1 labeling was also apparent in the neuropil of these regions, including prominent immunoreactivity consistent with juxtaparanodes of myelinated axons, similar to the staining pattern observed in the NTS (**Figure 3a-d**).

**Figure 3.**
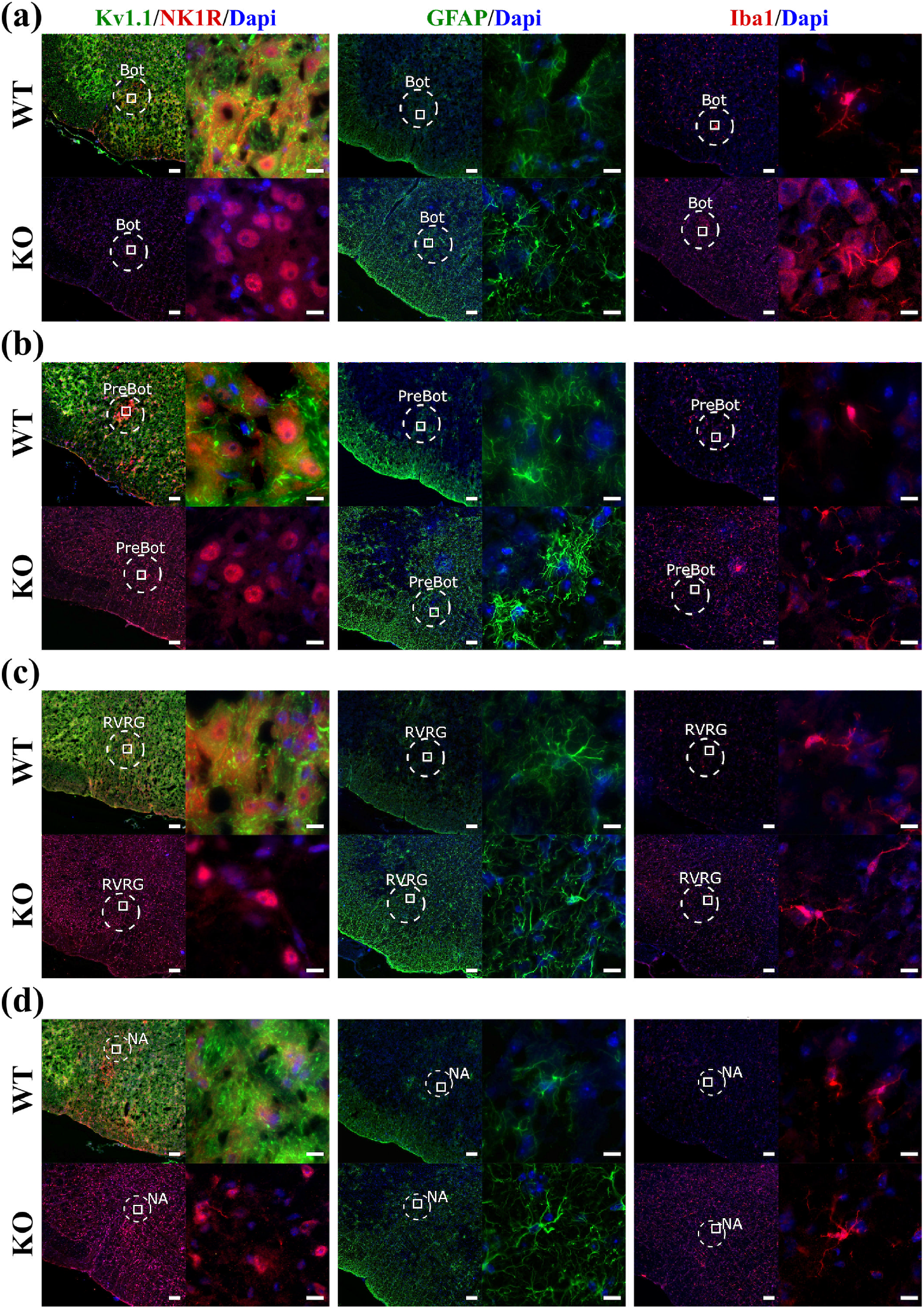
Kv1.1, GFAP, and Iba1 immunoreactivity in the ventral respiratory column and nucleus ambiguus (NA) of wild–type (WT) and *Kcna1^−/−^* (KO) mice. The structures shown include the (a) Bötzinger complex (Bot), (b) preBötzinger complex (preBot), (c) rostral ventral respiratory group (RVRG), and (d) NA. For each condition, the left panel shows a lower magnification image (scale bar = 100 μm) and the right panel shows a higher magnification image (scale bar = 10 μm) corresponding to the boxed region in the left panel. The dotted lines represent the boundary of the respective brain region being examined. DAPI staining was used to label the nuclei of cells. NK1R immunoreactivity, which is present in the neurons of the ventral respiratory column, was used to aid positive identification of the cells of the various nuclei.

The dorsolateral and ventrolateral pons are critical mediators of respiratory responses in the Hering–Breuer mechanoreflex and carotid chemoreflex pathways (Song and Poon 2004). These structures are connected reciprocally with one another and with medullary nucleus tractus solitarius (NTS) and VRG. Pontomedullary signal processing plays an important role in respiratory rhythm modulation and control of breathing function (Song and Poon 2004). The pontine respiratory group encompasses the parabrachial and Kölliker– Fuse nuclei within the dorsolateral pons, consisting of several types of neurons that modulate respiration (Ezure and Tanaka 2006). Forkhead box protein 2 (FoxP2), a transcription factor, is a highly expressed protein in parabrachial and Kölliker fuse nuclei (Geerling *et al*. 2011). Thus, Foxp2 immunoreactivity was used as a marker to identify the pontine respiratory group. Moderate Kv1.1 staining was observed in the neuropil and fibers tracts passing through the parabrachial and Kölliker–Fuse nuclei (**Figure 4a,b**). Kv1.1 immunoreactivity was observed in somata and neuronal processes of both nuclei.

**Figure 4.**
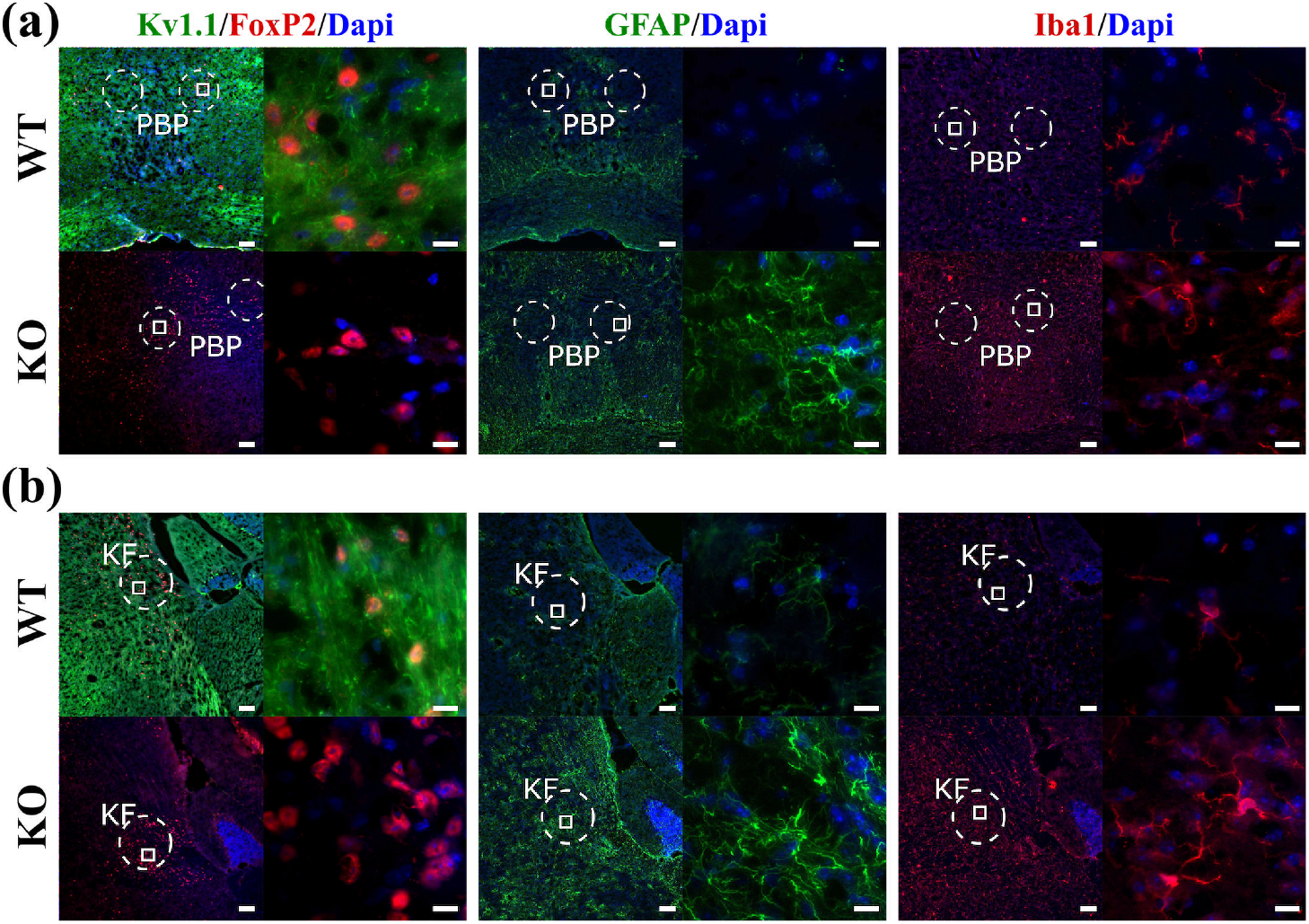
Kv1.1, GFAP, and Iba1 immunoreactivity in the pontine respiratory group of wild–type (WT) and *Kcna1^−/−^* (KO) mice. The structures shown include the in (a) parabrachial nucleus (PBP) and (b) Kölliker fuse nucleus (KF). For each condition, the left panel shows a lower magnification image (scale bar = 100 μm) and the right panel shows a higher magnification image (scale bar = 10 μm) corresponding to the boxed region in the left panel. The dotted lines represent the boundary of the respective brain region being examined. DAPI staining was used to label the nuclei of cells. FoxP2 immunoreactivity, which is present in the neurons of PBP and KF, was used to aid positive identification of these nuclei.

The retrotrapezoid nucleus (RTN), located in the rostral medulla, contains neurons that fire in response to increases in local pCO2 and that innervate the regions of the brainstem that contain the respiratory pattern generator (Guyenet *et al*. 2012). Strong Kv1.1 staining was observed in neuropil and juxtaparanodal-like puncta of the RTN where Kv1.1 may play a role in chemoreception (**Figure 5a**).

**Figure 5.**
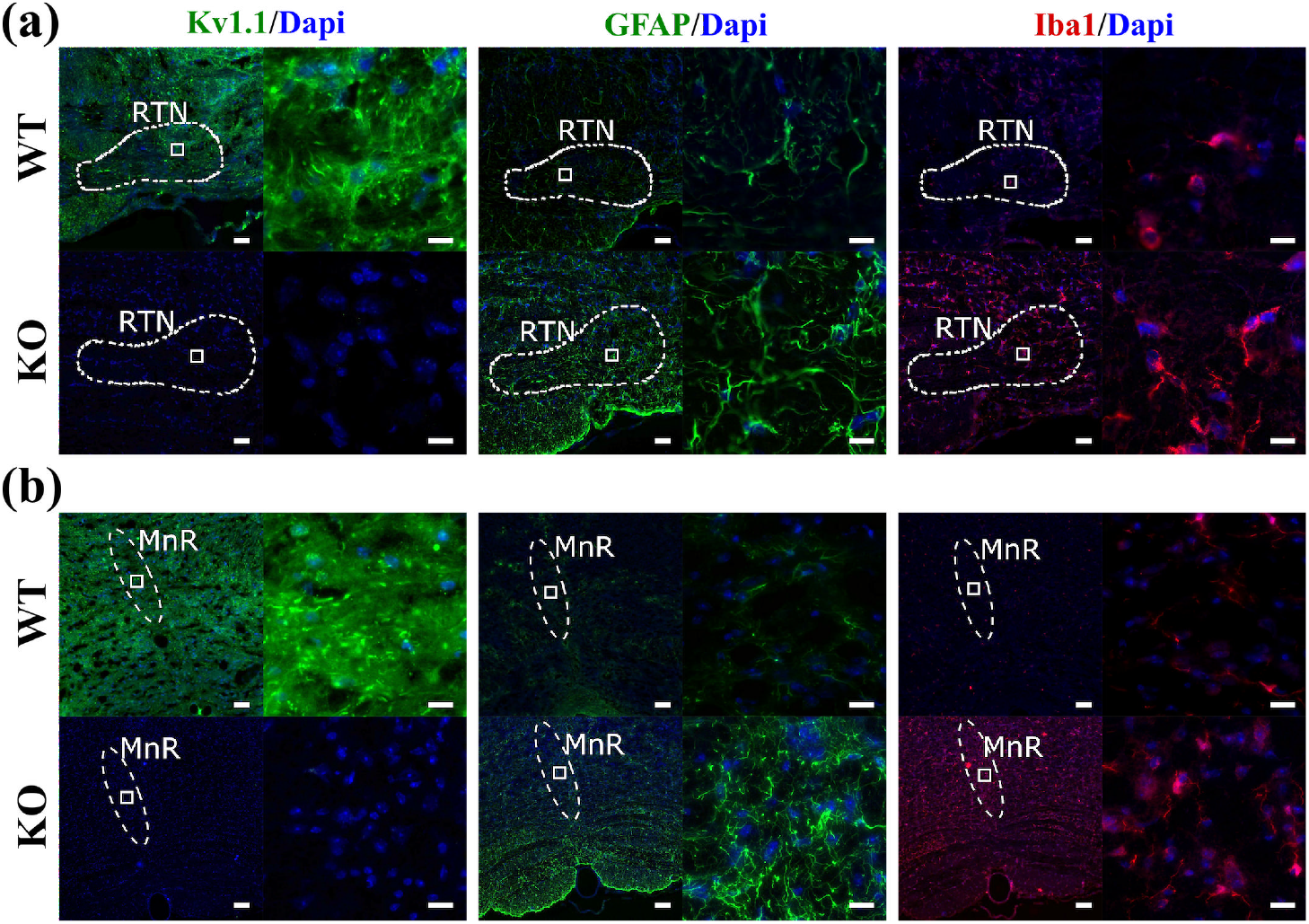
Kv1.1, GFAP, and Iba1 immunoreactivity in chemosensory brain regions in wild–type (WT) and *Kcna1^−/−^* (KO) mice. The structures shown include the (a) retrotrapezoid nucleus (RTN) and (b) median raphe nucleus (MnR). For each condition, the left panel shows a lower magnification image (scale bar = 100 μm) and the right panel shows a higher magnification image (scale bar = 10 μm) corresponding to the boxed region in the left panel. The dotted lines represent the boundary of the respective brain region being examined. DAPI staining was used to label the nuclei of cells.

Serotonergic neurons are important regulators of respiration and arousal and serotonergic signaling has been implicated in risk of sudden death in epilepsy (Richerson and Buchanan 2011; Hilaire *et al*. 2010). The raphe nuclei in the brainstem are the primary source of serotonergic neurons (Richerson and Buchanan 2011). High-density Kv1.1 immunoreactivity was observed in somata, neuropil, and juxtaparanodal-like puncta of the median raphae (**Figure 5b**).

### 3.2 Identification of astrogliosis

Seizures can injure the brain leading to reactive astrogliosis, which is defined by increased proliferation of astrocytes and characteristic changes in astrocytic morphology indicative of their reactive state, including hypertrophied cell bodies with abnormally long, extended processes. Thus, the presence of reactive astrocytes is an indicator of seizure-related brain damage that could lead to altered function (Sofroniew 2009; Patel *et al*. 2019). Using glial fibrillary acidic protein (GFAP) as a marker for astrocytes, we compared GFAP immunoreactivity in cardiorespiratory brain regions of adult WT (n=5) and KO (n=4) mouse brains to determine whether chronic seizures due to Kv1.1 deficiency induces astrogliosis. Overall, WT brains exhibited sparse GFAP immunoreactivity in all regions examined except for the VRC, where high densities of astrocytes are normally present, as previously reported (**Figures 1-6; Table 1)**(Grass *et al*. 2004; Sheikhbahaei *et al*. 2018). However, the GFAP-positive cells of the WT brains typically did not exhibit hypertrophic morphological changes indicative of reactive gliosis. In contrast, all KO brains showed extensive GFAP immunoreactivity throughout all cardiorespiratory brain regions examined, including a higher density of GFAP immunoreactive cells and an abundance of cells with enlarged somata and long processes consistent with reactive gliosis (**Figures 1-6)**. As summarized in **Table 1**, the density of GFAP immunoreactivity in KO brains relative to WT brains was highest in the basolateral amygdala, pontine respiratory group (parabrachial nucleus, Kölliker Fuse nucleus), and the median raphe nucleus. Moderate GFAP immunoreactivity was present in the NTS, 10N, 12N, and RTN of KO mice. The nuclei of the VRC also exhibited significant GFAP immunoreactivity but the increases were relatively lower compared to WT than the other cardiorespiratory brain regions (**Figures 1-6**).

### 3.3 Identification of microgliosis

Microglia are resident immune cells of the central nervous system constituting the first line of defense against pathogens or injury (Nimmerjahn *et al*. 2005). Similar to astrocytes, microglia proliferate and exhibit morphological changes in response to injury or infection leading to microgliosis (Fernández-Arjona *et al*. 2017; Torres-Platas *et al*. 2014). Using Iba1 (ionized calcium binding adaptor molecule-1) immunoreactivity as a specific marker of microglia, we compared microglial density between the brains of adult WT and KO mice to determine whether chronic seizures due to Kv1.1 deficiency induce microgliosis. In WT brains, Iba1-positive cells were present in all cardiorespiratory centers examined but they were relatively rare and low in abundance compared to KO mice (**Figures 1-6**). Conversely, in all KO mice, Iba-1 immunoreactivity appeared more abundant overall than in WT for all areas examined (**Figures 1-6**). As summarized in **Table 1**, the approximate densities of Iba1 immunoreactivity in KO brains (relative to WT) were highest in basolateral amygdala, Bötzinger complex, and median raphe nucleus, whereas moderate Iba1 immunoreactivity was present in 12N, preBötzinger complex, the Kölliker fuse nucleus and RTN. The RVRG, NA and parabrachial nucleus exhibited less Iba1 immunoreactivity.

## 4. DISCUSSION

This study provides the first description of Kv1.1 protein expression in the cardiorespiratory centers of normal mouse brain. Differential Kv1.1 immunoreactivity was detected in the main cardiorespiratory nuclei with the highest density of immunoreactivity observed in the NTS, hypoglossal nucleus, Botzinger nucleus, nucleus ambiguus, RTN, and median raphe nuclei. At the subcellular level, Kv1.1 immunoreactivity was evident in some cell bodies, but neuropil and axonal juxtaparanodal-like structures exhibited more prominent immunoreactivity. In *Kcna1^−/−^* mice, the presence of chronic seizures was associated with greater GFAP and Iba1 immunoreactivity, consistent with astrogliosis and microgliosis, respectively, which could impair brain function. Thus, the presence of Kv1.1 immunoreactivity in cardiorespiratory brain centers suggests that Kv1.1-containing channels could contribute to intrinsic control of breathing and that absence of Kv1.1 in *Kcna1^−/−^* mice could result in seizure-induced brain damage in the neurocircuits for respiratory function.

Any of the cardiorespiratory brain regions examined in this work could potentially contribute to the basal and seizure-associated cardiorespiratory phenotypes in *Kcna1^−/−^* mice, but one structure that is of special interest is the amygdala. Although not part of the canonical brainstem cardiorespiratory circuitry, limbic structures including the amygdala can exert powerful influence over respiration, especially during seizures (Dlouhy *et al*. 2015; Nobis *et al*. 2019; Lacuey *et al*. 2017; Applegate *et al*. 1983; Kaada and Jasper 1952; Harper *et al*. 1984). In patients, the onset of apneas is strongly correlated with the spread of seizures to the amygdala or with electrical stimulation of the amygdala (Dlouhy *et al*. 2015; Nobis *et al*. 2019; Lacuey *et al*. 2017). Stimulation of the hippocampus, another limbic structure which strongly expresses Kv1.1, can also evoke apneas in epilepsy patients (Lacuey *et al*. 2017; Wang *et al*. 1994). In *Kcna1^−/−^* mice, Fos immunostaining reveals that spontaneous seizures activate hippocampal and amygdalar circuits, which could contribute to the occurrence of ictal respiratory dysfunction in these animals (Gautier and Glasscock 2015; Dhaibar *et al*. 2019). The ability of the amygdala to influence respiration is likely due to its extensive projections to brainstem respiratory neurons, including connections with the respiratory rhythm generating pre-Botzinger complex (Yang *et al*. 2020; Price and Amaral 1981; Hopkins 1975). Thus, the absence of Kv1.1 could lead to neuronal excitability defects in the amygdala that alter respiration, and excessive activity due to seizures in *Kcna1^−/−^* mice could evoke downstream breathing dysfunction.

The brains of *Kcna1^−/−^* mice exhibited extensive activation of astrocytes and microglia throughout cardiorespiratory centers, indicative of brain injury sustained by the occurrence of repeated spontaneous seizures. Glial activation, or reactive gliosis, occurs in response to central nervous system insults such as seizures, and it is considered an integral part of epilepsy histopathology (Patel *et al*. 2019). The hallmarks of reactive gliosis include hypertrophy of cell bodies and processes and upregulated expression of GFAP and Iba1 in astrocytes and microglia, respectively (Patel *et al*. 2019). Reactive astrocytes increase neuronal hyperexcitability because they are unable to clear extracellular K^+^ and glutamate, resulting in reduced GABAergic inhibition of neurons (Patel *et al*. 2019; Robel and Sontheimer 2016). Reactive microglia also release proinflammatory cytokines which can lead to neuronal hyperexcitability and neurodegeneration that exacerbates epilepsy (Victor and Tsirka 2020; Hiragi *et al*. 2018). Previous studies have found varying degrees of astrogliosis in the hippocampus and hypothalamus of *Kcna1^−/−^* mice depending upon seizure severity, duration, and frequency (Wenzel *et al*. 2007; Roundtree *et al*. 2016). In addition, extensive seizure-related neurodegeneration and cell loss have been observed in the hippocampus and amygdala of *Kcna1^−/−^* mice, as well as in neocortex, piriform cortex and thalamus (Wenzel *et al*. 2007). Here, our study reveals that the extent of gliosis in *Kcna1^−/−^* mice extends beyond forebrain and limbic structures to include brainstem cardiorespiratory centers. Increased gliosis in brainstem respiratory control centers has also been observed in victims of sudden infant death syndrome (SIDS) (Takashima *et al*. 1978; Bruce and Becker 1991). Thus, the gliosis in the cardiorespiratory neurocircuits of *Kcna1^−/−^* mice observed here suggests that these brain networks are recruited during spontaneous seizures leading to pathological changes that could alter neuronal excitability and impair cardiorespiratory function.

This study provides an initial survey of Kv1.1 expression and gliosis in cardiorespiratory brain centers as a starting point for more detailed studies. However, the study contains several limitations that can guide these studies. First, Kv1.1 immunoreactivity was strongest for neuropil and fiber tracts rather than cell bodies, hindering the ability to unambiguously assign Kv1.1 expression to particular neuronal populations. For example, it is possible that the Kv1.1 immunoreactivity could be associated with axons or terminals that belong to different nearby structures that are synapsing on the neurons of interest. Future studies with tracer molecules and higher-resolution imaging could help distinguish the precise neuronal origin of Kv1.1-immunoreactive axons. More in-depth molecular characterization will also be required to determine the functional nature of the Kv1.1 immunoreactivity identified in this study (e.g., excitatory vs. inhibitory neurons; axonal vs. dendritic vs. synaptic terminal). Similarly, reactive astrocytes and microglia also exhibit distinct functional types (such as A1 vs. A2 for astrocytes or M1 vs. M2 for microglia) that are distinguishable by their specific molecular expression changes (Cherry *et al*. 2014; Li *et al*. 2019); discrimination of subtype-specific gliosis was beyond the scope of this study. More quantitative studies will be needed to derive exact counts of the extent of Kv1.1 expression in WT brain and gliosis in KO mice. Finally, cellular electrophysiological approaches will be required to determine the precise functional roles of Kv1.1 in the neural circuitry of the various cardiorespiratory brain regions.

In summary, this work describes widespread differential Kv1.1 expression in brain cardiorespiratory centers, which strengthens the hypothesis that Kv1.1 plays an important role in the intrinsic control of respiration. In *Kcna1^−/−^* mice, the presence of extensive gliosis in cardiorespiratory networks suggests the absence of Kv1.1 is associated with, if not causative for, seizure-related remodeling and damage. Thus, *Kcna1* gene deletion may lead to respiratory dysfunction either directly by inducing hyperexcitability in Kv1.1-deficient respiratory neurons, or indirectly by causing accumulation of seizure-related damage in respiratory brain networks.

## ACKNOWLEDGEMENTS

The study was supported by grants from the National Institutes of Health (NS100954 and NS099188 to EG) and by an Ike Muslow pre-doctoral fellowship from the Louisiana State University Health Sciences Center-Shreveport to H.D.

## AUTHOR CONTRIBUTIONS

HD and EG conceived and designed the study and wrote the manuscript. HD performed the experiments and analyzed the data. KAH assisted with image acquisition, data analysis and interpretation, and manuscript editing.

